# An Alternative, Cas12a-based CRISPR Interference System for Mycobacteria

**DOI:** 10.1101/2021.01.25.427309

**Authors:** Neil Fleck, Christoph Grundner

## Abstract

The introduction of CRISPR interference (CRISPRi) has made gene repression in mycobacteria much more efficient, but technical challenges of the prototypical Cas9-based platform, for example in multigene regulation, remain. Here, we introduce an alternative CRSPRi platform that uses the minimal Cas12a enzyme in combination with synthetic CRISPR arrays. This system is simple, tunable, and can regulate multiple genes simultaneously, providing a new tool to probe higher-order genetic interactions in mycobacteria including *Mycobacterium tuberculosis* (*Mtb*).

## INTRODUCTION

The adaptive bacterial immune systems based on clustered regularly interspaced short palindromic repeats (CRISPR) and CRISPR-associated proteins (Cas) have transformed genetics and the ease with which they can be programmed has led to their wide use in gene editing in eukaryotes and prokaryotes (1). One application of the CRISPR/Cas system, CRISPR interference (CRISPRi), introduced a new way of gene regulation by co-expressing an inactive Cas9 nuclease with an engineered single guide RNA (sgRNA) that directs the inactive nuclease to a target gene where it blocks transcription rather than cleaves the DNA (2). The prototypical CRISPRi system is based on an inactive type 2-II Cas9 nuclease (dCas9) and has recently also been adapted for use in mycobacteria including *Mtb* (3-5), for which genetic manipulation has long been an experimental bottleneck. Despite the advantages of CRISPRi, however, knockdown efficiency can vary widely for different target genes, and simultaneous manipulation of more than one gene remains challenging. These limitations have hampered attempts to probe redundant genes, gene families, and higher-order genetic interactions.

Natural CRISPR systems are inherently multi-gene regulatory systems that can principally also be reprogrammed to regulate multiple genes at once. One current limitation of the Cas9-based system, however, is the relatively large size of the sgRNA that requires a crRNA and a tracrRNA that are typically fused to produce the >100 bp long sgRNA. Expression of multiple sgRNAs for multigene knockdown requires stepwise cloning of large individual transcriptional units for each sgRNA and shows mostly moderate knockdown efficiency of 2-3-fold for most genes (6). The natural diversity of CRISPR systems, however, may offer simpler solutions to CRISPRi in *Mtb*: Recently, the minimal type 2-V CRISPR enzyme Cas12a (previously Cpf1) has been described (7). Cas12a does not require a tracrRNA and combines pre-crRNA processing and interference functions in one enzyme (8). The combination of these biochemical functions makes Cas12a a stand-alone enzyme that at least in some bacteria only requires a synthetic CRISPR array for the generation of mature crRNAs and subsequent gene repression (9, 10) (Fig. 1). Here, we adapted this system for CRISPRi in mycobacteria, creating a simple, highly tunable, reversible, multigene regulation CRISPRi platform.

**Figure 1.**
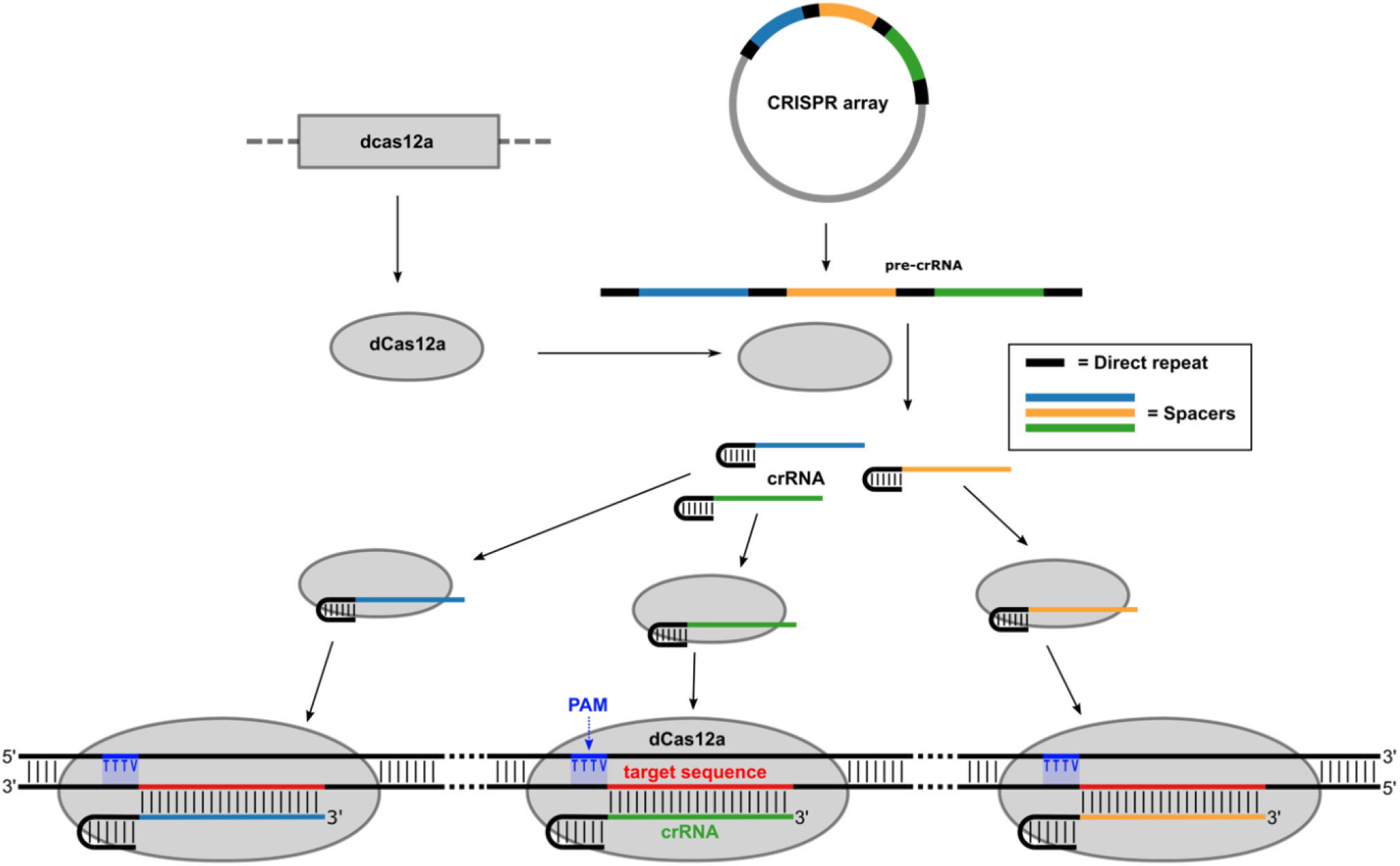
Schematic of dCas12a-based CRISPRi. Inducible, synthetic CRISPR arrays are transcribed into pre-crRNA and processed into mature crRNAs by dCas12a, which lacks DNA nuclease activity but retains RNA processing activity. The crRNAs direct dCas12a to the target DNA sequence(s), resulting in reduced transcription of one or multiple targets.

## RESULTS AND DISCUSSION

To test whether dCas12a in conjunction with a synthetic CRISPR array can be used for CRISPRi in mycobacteria, we stably introduced the gene for the inactive *Francisella novicida* Cas12a mutant Asp917Ala (dCas12a) into the Tweety recombination site of *Mycobacterium smegmatis* (*Msm*) strain mc^2^155 (11). The Asp917Ala mutant abrogates DNA cleavage activity but retains pre-CRISPR RNA processing activity (8). Initially, we could not detect the *Francisella*-derived dCas12a enzyme in *Msm* by Western blot, even after expression from a strong constitutive promoter. The *Francisella cas12a* sequence is AT rich (30% GC), while mycobacterial genomes are GC rich (*Mtb* 66%, *Msm* 67% GC). To test whether these differences limit expression, we tested a *Francisella cas12a* gene that was previously codon-optimized for human expression and has a GC content of 46%. This construct was readily expressed in *Msm* and *Mtb*. We next sought to test whether *Msm* expressing dCas12a can process synthetic CRISPR arrays into functional crRNAs and repress transcription of target genes. We first created the *Msm-luc-dCas12a* strain by integrating an ATc-inducible *dcas12a* into the Tweety recombination site, and integrating a constitutively expressed *luxCDABE* operon, which generates auto-luminescence in mycobacteria (12), into the L5 recombination site. We then targeted the template strand of the *luxCDABE* operon by episomally expressing synthetic CRISPR arrays. The CRISPR arrays contained the *Francisella* repeat sequence (GTCTAAGAACTTTAAATAATTTCTACTGTTGTAGAT) flanking each 22bp spacer sequence complementary to the luxCDABE promoter and coding sequence (Fig. 2A). Each target region on *luxCDABE* was selected to be immediately downstream from the protospacer adjacent motif (PAM) TTTV or TTN that licenses *Francisella* Cas12a binding to the target DNA (7, 13). The synthetic CRISPR arrays for production of pre-crRNA were expressed from an inducible mycobacterial expression plasmid under the control of ATc in the *Msm-luc-dCas12a* strain. After induction, we continuously measured luciferase activity for 16 hours. As controls, we included a strain expressing a non-targeting array (NTA) containing three 22bp spacer sequences without homology to *Msm* or *Mtb* and compared luminescence in each strain with and without ATc. We observed reduction of luminescence in all dCas12a strains carrying arrays with lux spacers when compared to the non-induced strains and the NTA. Reduction in luminescence was dependent on the number of spacers targeting the operon, from 15-fold with one targeting spacer (1T) to ∼45-fold with three (3T) and six spacers (6T), indicating that three spacers may be sufficient for maximal knockdown. Maximal knockdown was reached after 8-12 hours or 3-4 doubling times. To test the strand preference of dCas12a-mediated knockdown in mycobacteria, we next targeted the non-template strand of the lux promoter and sequences in *luxB* and *luxD* with a total of three spacers. Targeting the non-template strand reduced expression of lux genes, but to much lower degrees than the same number of spacers targeting the template strand (Fig. 2B), consistent with previous findings that the template strand is the main target for *Francisella* Cas12a (10, 13). Growth of *Msm* was marginally slowed by expression of dCas12a and the NTA (Fig. 2B). When we expressed dCas12a from a strong constitutive promoter, we observed baseline knockdown of about 10-fold even in the absence of ATc (data not shown), indicating that even small amounts of leaky expression of the array can lead to knockdown. Strains expressing both the array and dCas12a from ATc inducible promoters allowed for a wider range of induction and were used for this study. To explore the ATc dose response of knock down, we tested the effect of a range of ATc concentrations on luminescence (Fig 2C). Small concentrations of ATc as low as 4 ng/ml fully induced maximal knockdown, and lower concentrations induced partial knockdown. These data show that dCas12 can process pre-crRNA into mature crRNAs and repress endogenous genes in *Msm*.

**Figure 2.**
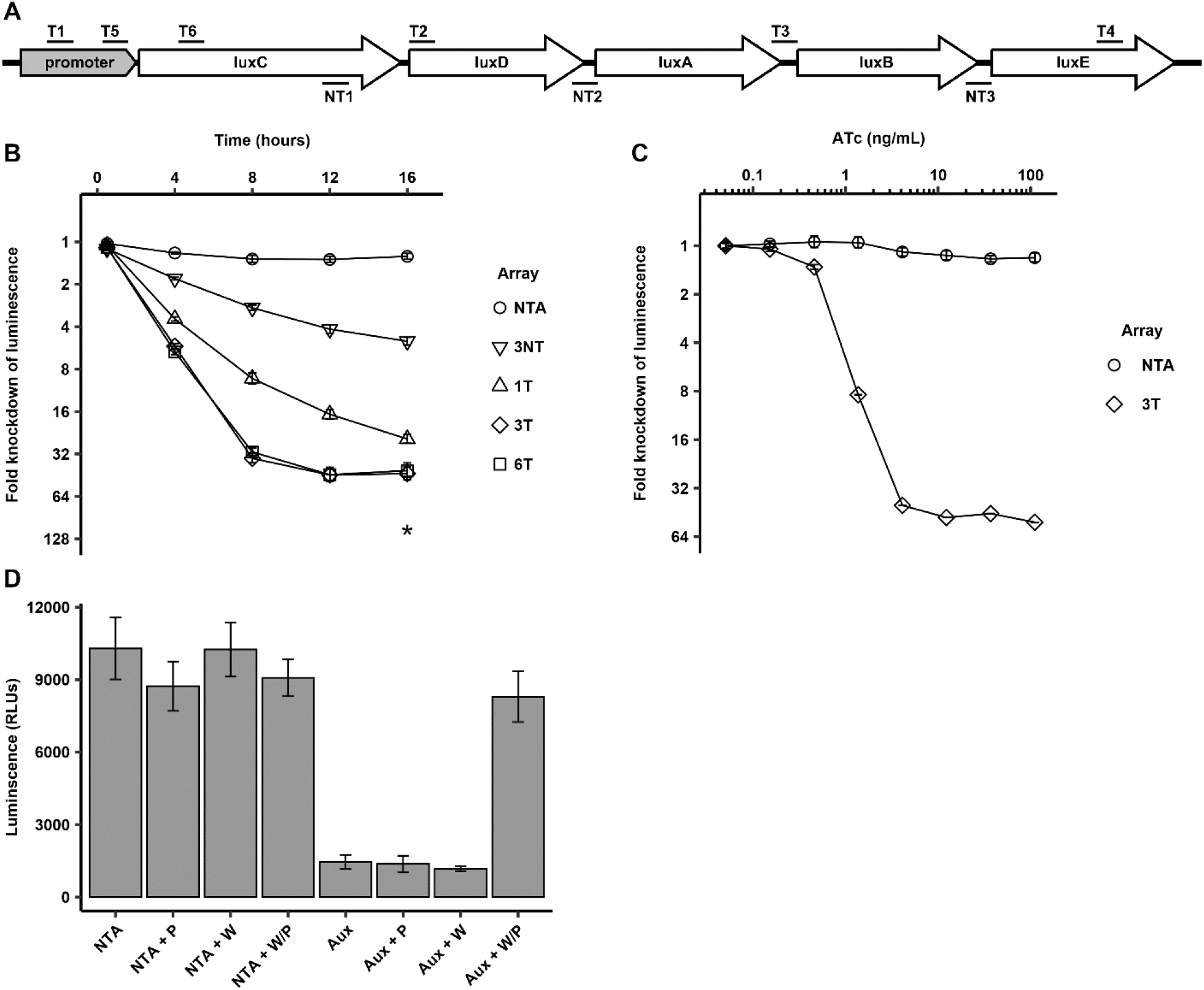
dCas12a-based CRISPRi in *Msm*. (A) Schematic of targeting locations of the spacers used to knock down the luxCDABE operon. T1-T6 mark the six spacers used to target the template (T) strand. NT1-3 mark the three spacers targeting the non-template (NT) strand. (B) Gene repression in *Msm*. Luminescence was knocked down in *Mtb-luc-dCas12a* strains by expressing synthetic CRISPR arrays with 1, 3, or 6 spacers targeting the luxCDABE operon or the non-targeting array. Knockdown of the lux operon was assessed by monitoring luciferase activity for 5 days post induction. Data are shown for each construct as the ratio of luminescence between the mock-induced samples and the ATc-induced samples for each construct. Array 1T contains spacer T1. Array 3T contains spacers T1, T3, and T4. Array 6T contains T1-T6, and array 3NT contains NT1, NT2, and NT3. NTA: Non-targeting array carrying three spacers against sequences not found in *Msm* or *Mtb*. Asterisk represents the assay’s limit of detection at the final timepoint. Error bars indicate standard error of six biological replicates. (C) ATc dose-response for tuning knockdown. *Msm-luc-dCas12a* strains expressing either the 3T array or NTA were grown to early log phase and induced with ATc from 0.07-100 ng/ml or mock-induced with DMSO, and luciferase knockdown was measured. Data are shown as the ratio of luminescence between the mock-induced strains and the ATc-induced strains after 16 hours. Error bars indicate standard error of three biological replicates. (D) Multigene repression in *Msm*. A synthetic array containing spacers targeting the essential Trp and Pro biosynthesis genes *trpD* and *proC* (Aux) was expressed in *Msm-luc-dCas12a*. Strains were grown in media supplemented with L-tryptophan (W) and L-proline (P), washed, diluted in media containing the indicated combinations of W and P supplementation, and growth was assessed after 12 hours by measuring luminescence. Only chemical supplementation of both amino acids recovers growth, indicating double knockdown of both *trpD* and *proC* genes. Error bars indicate standard error of five biological replicates.

Although the reduction of luciferase signal was likely due to the simultaneous repression of several lux genes, the common final readout for all lux genes (luminescence) in these experiments could not conclusively distinguish between single or multigene knockdown. To test for functional multigene knockdown, we next created a double auxotroph strain of *Msm* that carried a synthetic CRISPR array targeting two genes that are involved in the synthesis of the essential amino acids Trp (*trpD*) and Pro (*proC*). We introduced two spacers for each gene into the same CRISPR array. After induction of the array in *Msm* expressing dCas12a, the strain showed no growth in medium lacking either Trp and/or Pro (Fig. 2D). Complementation by the two amino acids fully restored growth to WT levels, whereas complementation with either one of the two amino acids did not rescue growth (Fig. 2D). These data show that both genes were effectively silenced and that dCas12a can target multiple endogenous genes.

We next tested whether dCas12a-mediated CRISPRi can be adapted to *Mtb*. We introduced synthetic CRISPR arrays containing spacers targeting the *luxCDABE* operon into *Mtb-luc-dCas12a*, an *Mtb* strain carrying dCas12a in the Tweety recombination site and the lux operon in the L5 recombination site. Induction of the arrays with ATc resulted in efficient reduction of luminescence (Fig. 3A). The degree of knockdown was dependent on the number of spacers targeting the operon, with one spacer resulting in 28-fold and three and six spacers resulting in 45-fold reduction of luminescence signal. Similar to *Msm*, three spacers were sufficient for maximal knockdown and knockdown was maximal after 3-4 doubling times. In contrast to *Msm*, targeting the non-template strand with three spacers did not reduce the luminescence signal. Expression of dCas12a and the NTA did not have an apparent effect on growth of *Mtb* (Fig. 3A). To test the reversibility of the system, we induced the *Mtb* strain carrying an array with three spacers targeting luc for four days. We next removed the inducer ATc by washout and resuspension in fresh medium. When ATc was added back to the cultures, the luciferase activity remained repressed. Without ATc, luciferase signal completely recovered after four days (Fig. 3B). These data show that the system is highly responsive and can be turned off within 3-4 doubling times by removing the inducer ATc. Next, we sought to test for multigene regulation in *Mtb*. We designed an array targeting the genes *pknH, fadD2, amiC, luxD*, and *rv0147* with 2 spacers for each gene-the 5x array. To detect knockdown of all targeted genes, determine the efficiency of knockdown, and to gauge potential off-target effects when expressing many spacers at once, we analyzed the knockdown strains by qRT-PCR. All five targeted genes were repressed, with between 3- and 14-fold reduction in mRNA (Fig. 3C). Transcript levels of the unrelated gene Rv0015c, which was not targeted by the array, was unchanged. These data show that at least five genes can readily be repressed by dCas12a simultaneously.

**Figure 3.**
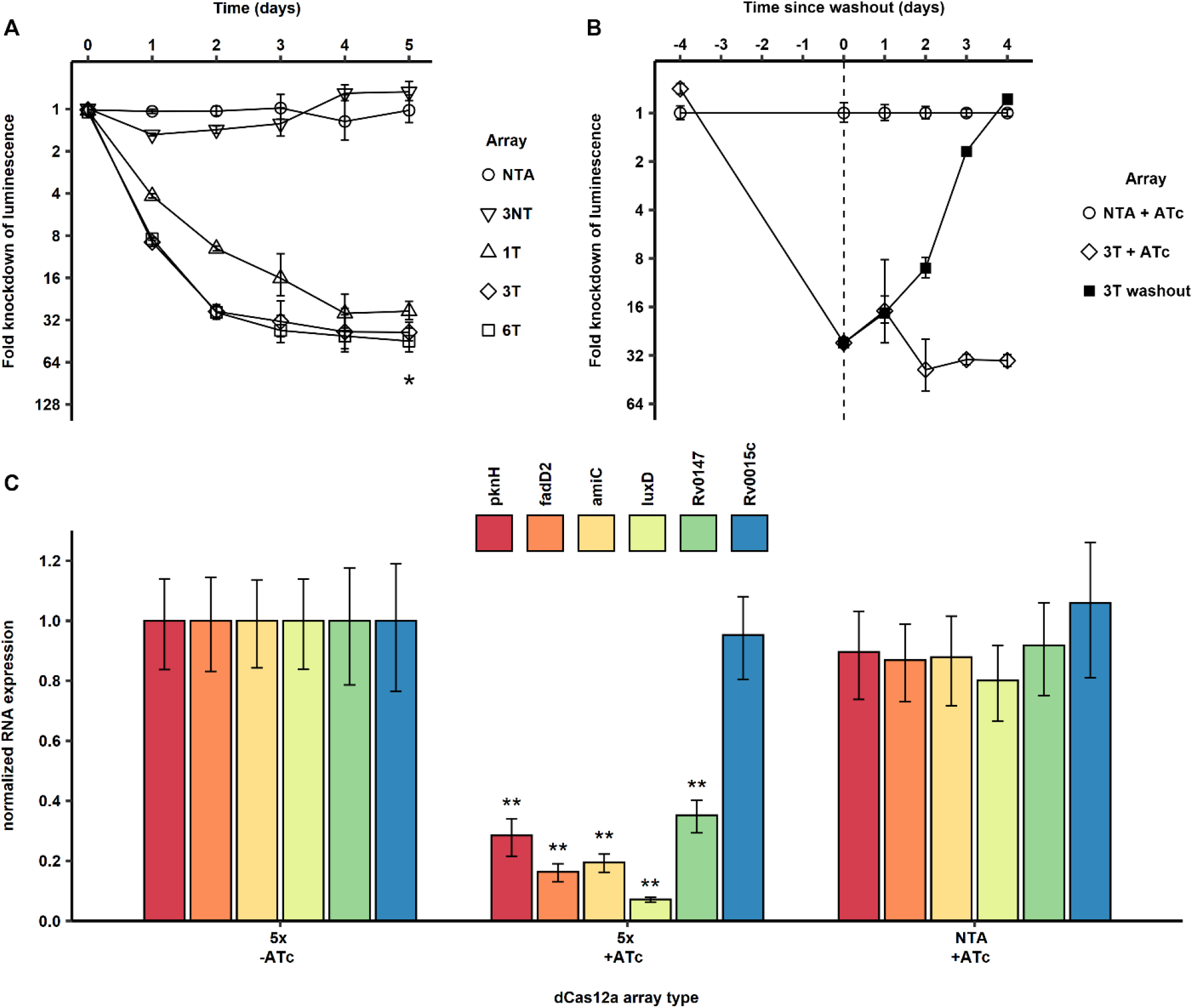
Cas12a-based CRISPRi in *Mtb*. (A) Gene repression in *Mtb*. Luminescence was knocked down in *Mtb-luc-dCas12a* strains by expressing synthetic CRISPR arrays with 1, 3, or 6 spacers targeting the luxCDABE operon or the non-targeting array. Knockdown of the lux operon was assessed by monitoring luciferase activity for 5 days after induction. Data are shown as the ratio of luminescence between the mock-induced samples and the ATc-induced samples for each construct. NTA: Non-targeting array carrying three spacers against sequences not found in*Mtb*. Asterisk represents the assay’s limit of detection at the final timepoint. Error bars indicate standard error of six biological replicates. (B) Reversibility of repression. After array induction and *luc* repression for four days, ATc was washed out. Luciferase signal remained repressed when ATc was added back, but completely recovered after four days without ATc. Luminescence was normalized to that of the NTA-carrying strain. Error bars indicate standard deviation of six biological replicates. (C) Multigene knockdown in *Mtb*. Five genes were targeted in *Mtb* by a single array (5x), and knockdown was measured by qRT-PCR. Rv0015c was not targeted by the 5x array expression and served as a control. NTA: Non-targeting array. Error bars indicate standard error of three biological replicates. Asterisks indicate p<0.001 compared to the mock-induced 5x strain.

Although the upper limit for the number of genes that can be repressed with this system remains to be determined, it might be higher than the five tested here: The natural *Francisella* CRISPR arrays contain between 5-26 spacers, perhaps indicating the natural upper limit for the system. The degree of knock down in the dCas12a-based system can be tuned by varying the concentration of the inducer ATc, varying the number of spacers targeting each gene, and by designing spacers with mismatches. For multigene knockdown, our data indicate a tradeoff between the efficiency of knockdown and the array size that limits the overall repression potential and that should be considered for optimal array design. The repetitive nature of the repeat sequences of the synthetic arrays also introduce cloning challenges, although we could readily obtain arrays with as many as 13 repeats by commercial gene synthesis.

Besides facile multigene regulation, Cas12a has additional advantages over Cas9-based CRISPRi systems. The much shorter repeat and spacer sequences (∼36bp and ∼22bp, respectively) and the single transcriptional unit required for any number of spacers make the Cas12a-based system more practical than the current Cas9-based system, which requires crRNAs of >100 bases with individual promoters. Limitations of Cas12a for CRISPRi, on the other hand, are the limited PAM sequences for the *Francisella* Cas12a in a GC-rich organism such as *Mtb*. However, the optimal PAM sequence TTTV (V=A/C/G) is found in 89% of *Mtb* coding sequences and is sufficient to target the *Mtb* coding sequence on average every 235 base pairs or 4 times per coding sequence, providing ample coverage. The PAM sequences for Cas12a derived from the core motif TTN are present in 98% of *Mtb* coding sequences and can be used at a small cost to knockdown efficiency (13, 14), and additional Cas12a orthologs with less stringent PAMs have recently been described (15), for example an engineered mutant of the Cas12a enzyme from *Acidaminococcus* (16) that recognizes the PAM sequence TYCV (Y=C/T), which is found in 98% of *Mtb* coding sequences.

Together, we introduce an alternative to the Cas9-based CRISPRi system for mycobacteria that allows for streamlined cloning, tunability, and efficient multigene regulation. This Cas12a-based system is versatile and the efficient multigene regulation will be particularly useful for the study of larger and redundant gene families and generally for the study of higher-order genetic interactions that have remained largely intractable.

**Figure 2 – Source Data 1**

Source data and statistical analysis descriptions for the luminescence knockdown experiments shown in Fig 2b, Fig 2c and Fig 2d (fig2-data1.xlsx)

**Figure 3 – Source Data 1**

Source data and statistical analysis descriptions for the luminescence knockdown and qRT-PCR experiments shown in Fig 3a, Fig 3b and Fig 3c (fig3-data1.xlsx)

## MATERIALS AND METHODS

### Media and growth conditions

*M. tuberculosis* and *M. smegmatis* were grown at 37°C in Middlebrook 7H9 broth or on 7H10 plates supplemented with 0.5% glycerol, 10% OADC, 0.05% Tween80 (broth only) with appropriate selective antibiotics and amino acids. Antibiotics were used at the following concentrations: ATc: 50 ng/ml (unless otherwise noted), Hyg: 100 ng/ml, Zeo: 25 µg/ml. L-tryptophan and L-proline were supplemented at 50 µg/ml.

### Array design and cloning

The pJEBTZ integrating plasmid was constructed by combining the Tweety-phage integrating backbone of pTTP1A (11), generously provided by the Hatfull lab, with the Zeocin selectable marker from psigE, the *E. coli* origin of replication from pJEB402-dCas10. Fn-Cpf1, with the nuclease deactivating mutation D917A and codon optimized for expression in humans, was sourced from pTE4999 (17) and inserted into the expression locus of pJEBTZ. The pJOBTZ vector was created by replacing the MOPS promoter in pJEBTZ with the TetR-regulated promoter p766 from pJR965 (4). The *Renilla* luciferase *LuxCDABE* operon was expressed under the control of the constitutive MOPS promoter on an L5-integrating plasmid containing a Kan-selectable marker.

The pNFCF vector was created by replacing the Uv15-Tet promoter in pDTCF (18) with the synthetic TetR-regulated promoter p766 from pJR965. To clone crRNA arrays into pNFCF, sequences flanked by overhangs for Gibson Assembly matching the vector insertion site (5’-CCGCATGCTTAATTAAGAAGGAGATATACAT-3’) – array sequence – (5’-GACTACAAGGATGACGACGACAAG-3’) were synthesized (Genscript) and obtained in a pUC57-Kan vector. The inserts were amplified using standard PrimeStarHS PCR chemistry with 55°C annealing temp and the forward primer (5’-CCGCATGCTTAATTAAGAAGGAGATATACAT-3’) and the reverse primer (5’-CTTGTCGTCGTCATCCTTGTAGTC-3’). The pNFCF vector was PCR linearized with forward (5’-ATGTATATCTCCTTCTTAATTAAGCATGCGG-3’) and reverse (5’-GACTACAAGGATGACGACGACAAG-3’) linearization primers, and a touchdown thermocycler program using 60°C to 55°C annealing temperatures. Array inserts and linearized pNFCF vector were gel-purified and then assembled via Gibson Assembly.

### Autoluminescence assays

Strains in log phase were grown for at least 2 doublings and diluted to OD_600_ of 0.005, and expression of the crRNA array and dCas12a was induced with ATc or mock-induced with DMSO. To assess knockdown, 100ul culture was dispensed into 96-well, white, flat-bottom plates, and luminescence was quantified using a PHERAstar plate reader (BMG Labtech), blanked against media. For *M. smegmatis*, cultures were kept inside the plate reader at 37°C for the duration of the experiment.

### Auxotroph supplementation assay

*M. smegmatis* cultures were grown to late log-phase in media supplemented with 50 μg/ml L-tryptophan and L-proline, then diluted to an OD_600_ of 0.4 and grown for 3 hours in the dark. Cultures were washed twice in non-supplemented media and diluted to an OD_600_ of 0.1 with ATc and Hyg to drive expression of and maintain selection for the CRISPRi system. Different combinations of amino acids were supplemented into the cultures, and growth was monitored after 12 hours at 37°C by measuring luminescence.

### qRT-PCR analysis of multigene knockdown

Liquid cultures of *Mtb-luc-dCas12a* were induced in early log phase with 50 ng/ml ATc or mock-induced with DMSO and grown for 4 days to mid-log phase. RNA was extracted in Trizol, purified, and cDNA was synthesized using the SuperScriptIV polymerase and random hexamer primers. mRNA expression levels for each gene were determined by qRT-PCR using SybrGreen iTaq chemistry and normalized to sigH mRNA expression and plotted as the ratio of mRNA levels (calculated as 2^-ΔΔCq^) in each strain relative to the mock-induced 5x strain.

## ACKNOWLEDGEMENTS

This work was supported by NIH grants R01AI117023, R21AI137571, and R03AI131223 and by a grant by the American Lung Association to C.G. We thank Jeremy Rock and Graham Hatfull and lab members for vectors and advice with cloning. Plasmid pTE4889 was a gift from Ervin Welker (Addgene plasmid # 88905).Plasmid pTTP1A was a gift from Graham Hatfull (Addgene plasmid # 91721)

## SUPPLEMENTARY DATA

**Table S1:**
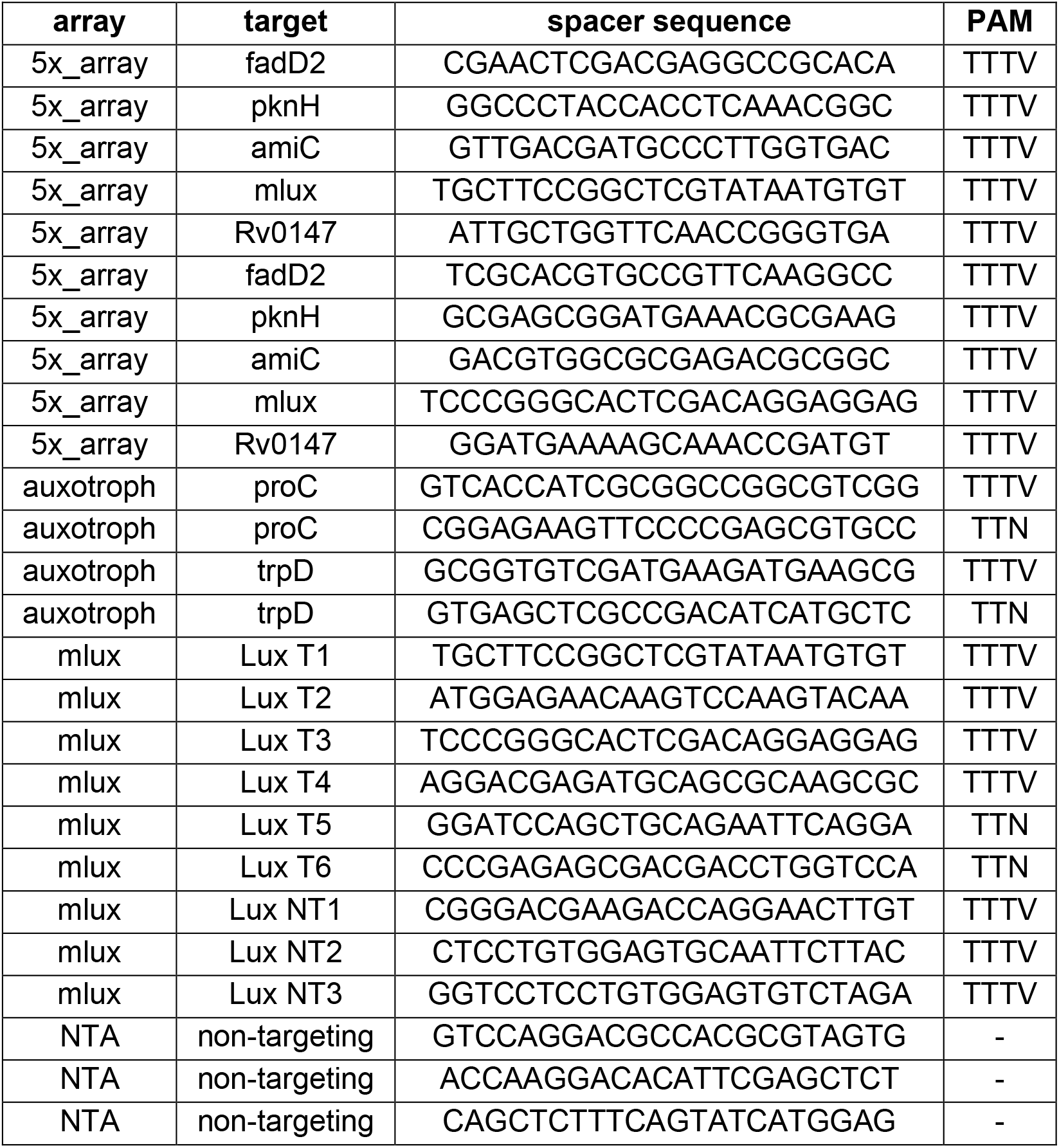
crRNA array spacers

**Table S2:** plasmids used in this study

| Plasmid | Relevant Features | Antibiotic resistance | Reference |
| --- | --- | --- | --- |
| pJOBTZ_hdcas12a | Plasmid for integrating ATc-inducible dcas12a (codon optimized for humans) into the Tweety integration site of Mycobacteria | Zeo | This study |
| pJEBTZ_hdcpf1 | Plasmid for integrating constitutively expressed dcas12a (codon optimized for humans) into the Tweety integration site of Mycobacteria | Zeo | This study |
| pmlux | Plasmid for integrating constitutively expressed luxCDABE operon into mycobacterial L5 site | Kan | Andreu N, et al. (12) |
| pNFCF_Cpf1_mlux1c | ATc inducible cas12a array containing one spacer targeting luxCDABE | Hyg | This study |
| pNFCF_Cpf1_mlux3c | ATc inducible cas12a array containing three spacers targeting luxCDABE | Hyg | This study |
| pNFCF_Cpf1_mlux3t | ATc inducible cas12a array containing three spacers targeting luxCDABE (non-template strand) | Hyg | This study |
| pNFCF_Cpf1_mlux6c | ATc inducible cas12a array containing six spacers targeting luxCDABE | Hyg | This study |
| pNFCF_Cpf1_NTA | ATc inducible cas12a array containing three randomly generated non-targeting spacers which have no homology to Mtb or Msm | Hyg | This study |
| pNFCF_Cpf1_PFAM0147 | ATc inducible cas12a array containing two spacers each for targeting pknH, fadD2, amiC, luxCDABE operon, and Rv0147 in Mtb | Hyg | This study |
| pNFCF_Cpf1_aux3 | ATc inducible cas12a array containing two spacers each for targeting proC and trpD in Msm | Hyg | This study |

